# UBQLN2 facilitates degradation of the retrotransposon protein PEG10 via UBE3A activity

**DOI:** 10.1101/2025.03.28.646022

**Authors:** Julia E. Roberts, Phuoc T. Huynh, Alexandra M. Whiteley

**Affiliations:** Department of Biochemistry, University of Colorado Boulder, Boulder CO 80309; Department of Molecular, Cellular, and Developmental Biology, University of Colorado Boulder, Boulder CO 80309

## Abstract

Ubiquilins are a family of extrinsic ubiquitin receptors that are thought to facilitate protein degradation by shuttling proteins to the proteasome. However, the defining characteristics of Ubiquilin clients, and the steps of Ubiquilin-mediated degradation, have been elusive. Previously, we showed Ubiquilin 2 (UBQLN2) regulates the proteasomal degradation of PEG10, a unique virus-like protein which comes in two forms: a gag protein which is not regulated by UBQLN2, and a gag-pol protein which is dependent on UBQLN2. Here, we refine the model of Ubiquilin activity through the UBQLN2-mediated degradation of PEG10. UBQLN2 binding did not ensure degradation, and was independent of client ubiquitination, though ubiquitination of key lysine residues was necessary for gag-pol proteolysis. Ubiquitination was dependent on the E3 ubiquitin ligase UBE3A, which was surprisingly unable to regulate gag-pol in the absence of UBQLN2. Together, we have established a stepwise model of UBQLN2-mediated degradation that represents a new perspective on Ubiquilin function.

**Graphical abstract: Working model of UBQLN2 and UBE3A-mediated degradation of PEG10 gag-pol.** In an apparent first step, UBQLN2 binds both PEG10 gag and gag-pol through STI1:gag-dominated interactions, independent of PEG10 ubiquitination. Either concurrently or immediately thereafter, UBQLN2 binds to UBE3A through interactions facilitated by UBA:AZUL domain binding, though other protein domains contribute to this interaction as well. UBE3A and other unknown E3 ligases contribute to the ubiquitination of gag-pol on lysine residues of both the gag and pol regions, which is necessary for proteasomal degradation. The proteasome also interacts with UBQLN2:UBE3A:PEG10, likely through UBL domain interactions with the regulatory cap. Inhibition of E1 ubiquitin activation through TAK243, UBE3A activity with siRNA, or proteasomal degradation with Bortezomib all interfere with UBQLN2-mediated PEG10 degradation.

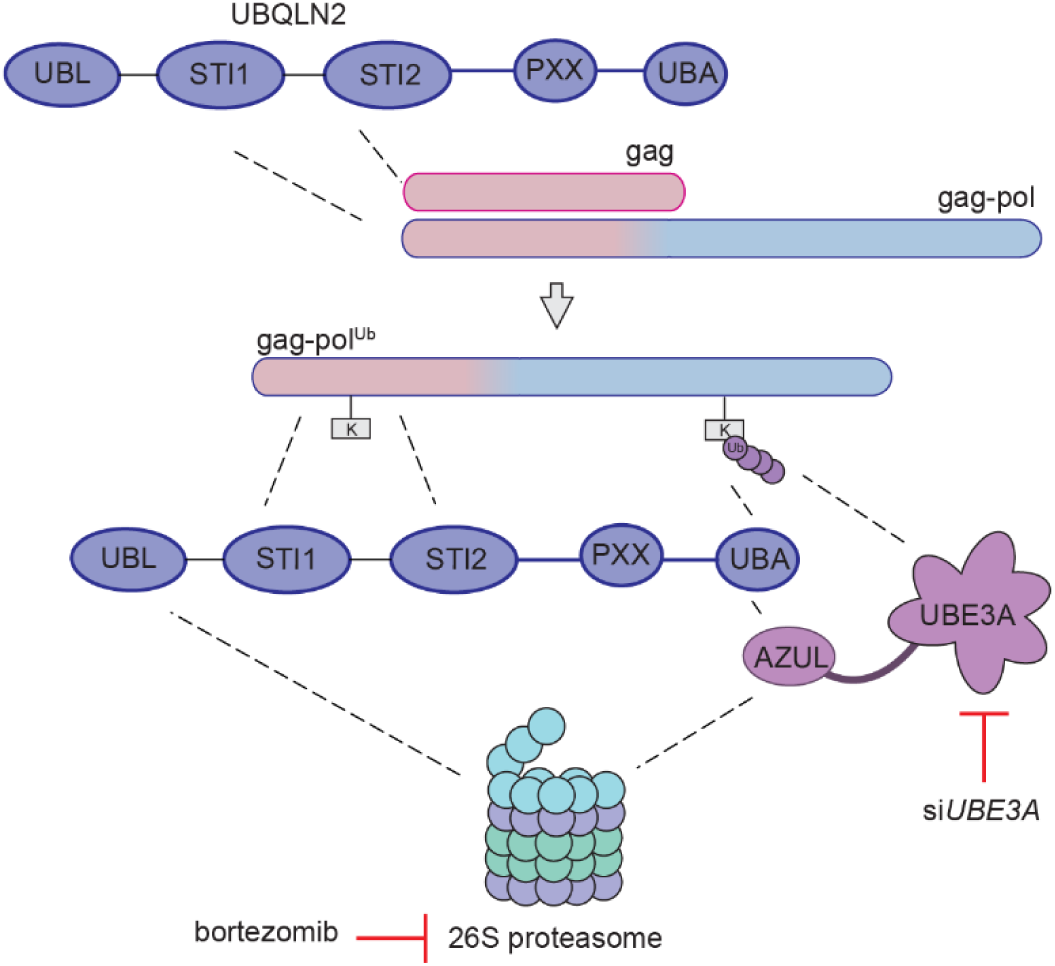

## Introduction

Protein degradation is essential to cellular and organismal health, and the majority of proteins within the cell are degraded by the Ubiquitin Proteasome System (UPS), where a cascade of enzymes mark proteins for degradation through the covalent attachment of ubiquitin chains at one or more lysine residues (Hershko *et al*., 1983; Hershko and Ciechanover, 1998; Nandi *et al*., 2006). Ubiquitinated proteins are then recognized at the regulatory cap of the proteasome via intrinsic ubiquitin receptors (Rpn10, Rpn13, and Rpn1), which facilitate docking and subsequent threading of proteins into the core for proteolytic cleavage (Finley, 2009; Lander *et al*., 2012).

While many proteins are degraded efficiently through the UPS, a subset of proteins rely on the assistance of extrinsic ubiquitin receptors, or ‘shuttle factors’. These receptors bind to the proteasome through a ubiquitin-like (UBL) domain (Elsasser *et al*., 2002; Chen *et al*., 2016; Buel *et al*., 2023) and bind to ubiquitin moieties through a ubiquitin associated (UBA) domain (Funakoshi *et al*., 2002; Kang *et al*., 2006; Harman and Monteiro, 2019). Ubiquilins (UBQLNs) are the largest mammalian family of extrinsic ubiquitin receptors, and additionally contain either one or two intervening STI1 domains thought to facilitate protein-protein interactions (Hjerpe *et al*., 2016; Suzuki and Kawahara, 2016; Zheng, Yang and Castañeda, 2020; Fry, Saladi and Clemons Jr, 2021; Onwunma *et al*., 2024), including those with clients (Kurlawala *et al*., 2017). There are five UBQLNs expressed in humans: UBQLNs 1, 2, 3, 4, and L, which all share the same general protein domain structure, but diverge in tissue expression (Lin *et al*., 2022). UBQLN1 and UBQLN4 are ubiquitously expressed, while UBQLN2 is enriched in muscle tissue and the nervous system, and UBQLN3 and UBQLNL are restricted to the testes (Marín, 2014). UBQLN2 has also gained recent attention as mutations cause a familial form of Amyotrophic Lateral Sclerosis (fALS) with frontotemporal dementia (FTD) (Deng *et al*., 2011; Wu *et al*., 2020; Lin *et al*., 2022).

An unresolved question within the Ubiquilin field has been the identity of proteins, called ‘clients’, which rely on Ubiquilin activity for degradation. Loss of the *UBQLN* ortholog *Dsk2* results in a proteome-wide accumulation of K48-linked ubiquitinated proteins (Funakoshi *et al*., 2002), which supports the notion that Dsk2 is a general facilitator of ubiquitinated protein degradation. However, mammalian UBQLNs may play a more complex role. Mammalian UBQLNs bind with nearly equal affinity to a variety of ubiquitin linkages, making K48-linked ubiquitination unlikely as the single client feature (Zhang, Raasi and Fushman, 2008). In addition, recent work has observed UBQLN-dependent proteasomal degradation of non-ubiquitinated proteins (Makaros *et al*., 2023).

In a model where ubiquitination is not sufficient to mark a protein for UBQLN-dependent degradation, considerable work has been done to identify classes of Ubiquilin clients of shared biophysical or functional properties. UBQLN2 has been linked to the degradation of aggregated proteins (Hjerpe *et al*., 2016), UBQLN4 has been linked to the degradation of nuclear MRE11 (Suzuki and Kawahara, 2016), and UBQLN1/UBQLN2 have been tied to the degradation (or protection, in some cases) of mislocalized mitochondrial proteins (Itakura *et al*., 2016; Whiteley *et al*., 2017). Considerable work has also tied Ubiquilins to the degradation of proteins involved in neurodegeneration, including presenilins (Harman and Monteiro, 2019), huntingtin (Chuang *et al*., 2016), TDP43 (Kim *et al*., 2009), and others (Zheng, Yang and Castañeda, 2020). Recently, consensus is emerging that mild hydrophobic stretches are a common theme of Ubiquilin clients (Itakura *et al*., 2016; Whiteley *et al*., 2017; Onwunma *et al*., 2024). However, this does not appear to be exclusive, as there are many reported clients without clear hydrophobic stretches (Zheng, Yang and Castañeda, 2020).

In addition to this unresolved question of client selection, the field lacks a clear chronological understanding of how Ubiquilin clients are degraded in a stepwise manner. While there is evidence Ubiquilins can facilitate degradation of both ubiquitinated (Zheng, Yang and Castañeda, 2020) and non-ubiquitinated proteins (Makaros *et al*., 2023), recent evidence points to a putative model in which ubiquitination of protein is instead facilitated by Ubiquilin interaction (Itakura *et al*., 2016; Onwunma *et al*., 2024), thereby facilitating either degradation or protection of proteins. Deep characterization using model client proteins can help clarify this new model of Ubiquilin function.

In this study, we used PEG10 as a reporter system for exploring the process of UBQLN2-mediated degradation. *Paternally expressed gene 10 (PEG10)* is a unique retrotransposon-derived gene which retains many structural features of retrotransposons, but has lost the enzymatic domains necessary for genetic duplication (Brandt, Veith and Volff, 2005). However, it has retained a -1 programmed ribosomal frameshift, which allows for the formation of two functionally unique protein products from a single mRNA, which we refer to as gag and gag-pol (Brandt, Veith and Volff, 2005; Clark *et al*., 2007; Lux *et al*., 2010). The PEG10 gag-pol protein is a genuine UBQLN2 client, and is an excellent client reporter (Whiteley *et al*., 2021): it is degraded in a proteasome- and UBQLN2-dependent fashion, and can be fused to a fluorescent protein without losing Ubiquilin dependence (Black *et al*., 2023). While gag-pol is a UBQLN2 client, gag is not, providing a unique opportunity to explore client identity and the chronology of Ubiquilin-mediated degradation (Black *et al*., 2023).

In this study, we determined that PEG10 gag-pol is degraded in a ubiquitin, proteasome, and UBQLN2-dependent fashion, and that key lysine residues are necessary for this degradation pathway. Ubiquitination was not necessary for UBQLN2 binding but was necessary for proteasomal degradation. Ubiquitination of PEG10 occurs via the E3 ligase UBE3A, which required the presence of UBQLN2 to regulate gag-pol abundance. In conclusion, we determined the chronology and context with which Ubiquilin-mediated degradation of PEG10 gag-pol occurs. Together, these findings help refine our understanding of Ubiquilin function and their mechanism of action, which will inform future studies of Ubiquilins, their clients, and E3 ligases.

## Results

### PEG10 gag-pol degradation is dependent on the ubiquitin proteasome system

While many reported Ubiquilin clients are destined for degradation through the UPS (Hjerpe *et al*., 2016; Itakura *et al*., 2016), some clients are degraded in a ubiquitin-independent pathway (Makaros *et al*., 2023), and others are degraded through autophagy (Rothenberg *et al*., 2010; Yun Lee, Arnott and Brown, 2013; Wu *et al*., 2020; Idera *et al*., 2023). To determine the pathway of degradation for PEG10, endogenous gag and gag-pol protein abundance was probed in WT HEK293 cells upon proteasome and E1 inhibition by bortezomib and TAK243, respectively. As expected, poly-ubiquitinated proteins increased in total cell lysate upon bortezomib treatment due to the inhibition of proteasomal degradation (**Figure 1A**). Conversely, poly-ubiquitinated proteins were decreased, while mono-ubiquitin accumulated, upon TAK243 treatment (**Figure 1A**). Treatment of cells with both TAK243 and bortezomib prevented the accumulation of poly-ubiquitinated protein (**Figure 1A**). Gag protein levels were not significantly impacted by either drug (**Figure 1A-B**) but gag-pol accumulated significantly upon either proteasome or E1 inhibition (**Figure 1A-B**), indicating that gag-pol abundance is uniquely regulated by ubiquitination and proteasomal degradation. When the molecular weight of gag-pol was carefully examined, high molecular weight signal (120-250 kDa) increased relative to tubulin expression only upon bortezomib treatment (**Supplementary Figure 1A**), suggesting an accumulation of poly-ubiquitinated gag-pol protein. When low and high molecular weight gag-pol proteins were compared, their ratio did not change upon bortezomib treatment (**Supplementary Figure 1B**), which may reflect inhibition of PEG10 ubiquitination due to proteasome inhibition. In comparison, the ratio of high molecular weight PEG10 decreased upon treatment with the E1 inhibitor TAK243, and with combined treatment (**Supplementary Figure 1B**), further suggesting that this high molecular weight species includes ubiquitinated protein.

**Figure 1:**
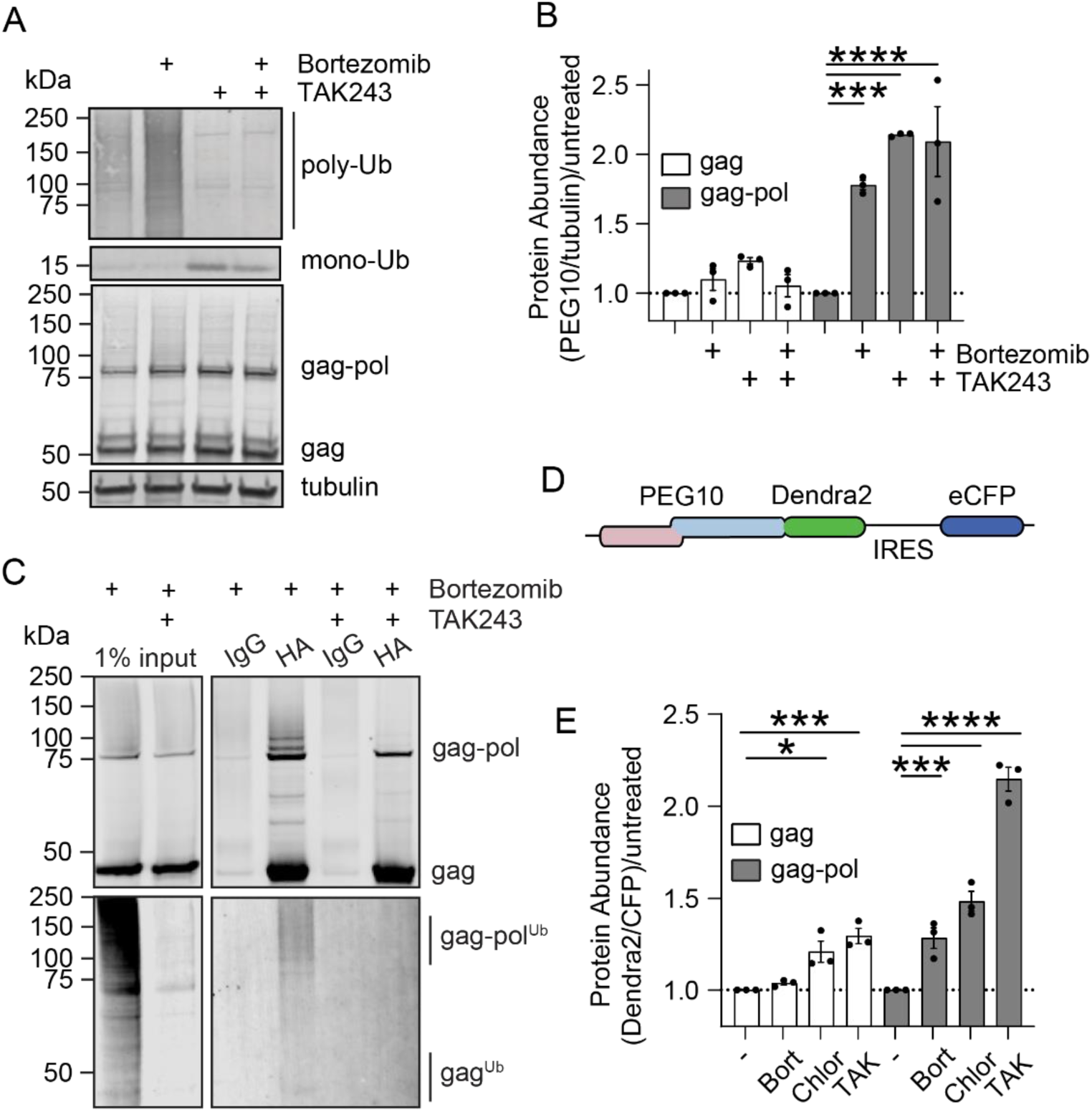
PEG10 gag-pol degradation is dependent on the Ubiquitin Proteasome System (UPS). **A)** Representative western blot of WT HEK293 whole cell lysate. Cells were treated with drug for 16 hours prior to harvest. Poly-ubiquitin and mono-ubiquitin were probed with pan-Ub antibody (P4D1); PEG10 was probed with polyclonal PEG10 antibody. Shown is one of three representative experiments. **B)** Quantification of data shown in (A), where samples were normalized to the abundance of each protein in untreated cells for each individual experiment. Statistics were determined by two-way-ANOVA with multiple comparisons, n=3. **C)** Representative western blot analysis following immunoprecipitation using HA-tag antibody or IgG control from whole cell lysate of cells stably expressing HA-gag-pol and transiently transfected with 3xFLAG-Ubiquitin. Cells were treated with drug for 16 hours prior to harvest and IP. Western blot was probed either against HA (top) or FLAG (bottom). Shown is one of two representative experiments. **D)** Schematic of Dendra2 reporter. Gag region of PEG10 is shown in pink, and pol region is shown in blue. Dendra2 is fused at the C-terminus to gag-pol and will only be visible when the ribosome has successfully frameshifted. IRES: internal ribosome entry site; eCFP: enhanced CFP. **E)** Client accumulation assay performed using gag and gag-pol, with and without drug treatment. Cells were treated with the drugs indicated for 16 hours prior to harvest and flow cytometry. Like (B), samples were normalized to the abundance of each protein in untreated (-) cells for each individual experiment. Statistics were determined by one-way-ANOVA with multiple comparisons, n=3. For (B,E), data are shown as mean ± SEM.

Accumulation of gag-pol upon proteasome and E1 inhibition suggested that the protein is ubiquitinated as a prerequisite for degradation by the proteasome. To monitor ubiquitination of PEG10 more directly, a 3xFLAG-Ubiquitin (3xFLAG-Ub) plasmid was transfected into a cell line stably expressing HA-tagged PEG10. Cells were treated with bortezomib to enhance PEG10 yield for immunoprecipitation (IP) in the presence or absence of TAK243. When probed against HA, a ladder of high molecular weight gag-pol was visible with bortezomib treatment, which disappeared upon co-treatment with TAK243 (**Figure 1C**). Similarly, a smear of 3xFLAG-Ub at the same molecular weight was visible in HA-precipitated samples (**Figure 1C** and **Supplementary Figure 1C**) which disappeared upon TAK243 treatment. In contrast, no obvious ubiquitin signal was visible at or near the molecular weight for gag (**Figure 1C**).

Similar results were found using a fluorescent reporter of PEG10 accumulation. In brief, fusion constructs are generated between proteins of interest and a Dendra2 fluorescent protein, followed by an IRES-CFP cassette (**Figure 1D**). Cells were transfected with reporter constructs, and flow cytometry was performed 48 hours later. The ratio of Dendra2/CFP provides transfection-controlled protein abundance values with single-cell granularity. As gag abundance was not impacted by proteasome or E1 inhibition in our previous experiment, an inhibitor of autophagy, chloroquine, was added. In this assay, gag protein accumulated to a minor amount upon inhibition of autophagy and ubiquitination (**Figure 1E**). In contrast, PEG10 gag-pol accumulated upon inhibition of the proteasome, autophagy, and ubiquitination, which more than doubled upon incubation of cells with TAK243 (**Figure 1E**).

### Proteasomal degradation of PEG10 gag-pol is Ubiquilin-dependent

To determine how Ubiquilins contribute to these protein degradation pathways, the same assays were performed in a triple knockout Ubiquilin HEK293 cell line lacking Ubiquilins 1, 2, and 4 (TKO). First, endogenous levels of PEG10 were quantified by western blot in TKO cells after proteasome or E1 inhibition (**Figure 2A**). In contrast to WT HEK cells, TKO cells showed a significant increase in gag protein upon treatment with TAK243, but not with bortezomib alone (**Figure 2B**), implying abundance was regulated by ubiquitination but not by proteasomal degradation. Similarly, gag-pol was not regulated by the proteasome but was impacted by E1 inhibition in TKO cells (**Figure 2B**). Closer examination of high molecular weight products showed that TKO cells demonstrate a similar magnitude of increase compared to WT cells upon bortezomib treatment (**Supplementary Figure 1A, Supplementary Figure 2A**). However, when comparing the ratio of 80 kDa protein to higher molecular weight products, bortezomib treatment caused a doubling of the ratio of high molecular weight species relative to the 80 kDa gag-pol protein (**Supplementary Figure 2B**). In comparison, WT cells treated with bortezomib did not show a relative change in the ratio of PEG10 proteins upon bortezomib treatment (**Supplementary Figure 1B**). From this, we conclude (1) at the steady state, TKO cells had a disproportionate amount of 80 kDa gag-pol protein, which is likely not ubiquitinated, and which is insensitive to proteasome inhibition in TKO cells. We also conclude that (2) the small proportion of ubiquitinated gag-pol in TKO cells is capable of being degraded by the proteasome, as it accumulates upon bortezomib treatment. Western blot results were validated by client accumulation assay which showed gag-pol abundance did not change upon proteasomal inhibition in TKO cells (**Figure 2C**), in contrast with WT cells (**Figure 1E**).

**Figure 2:**
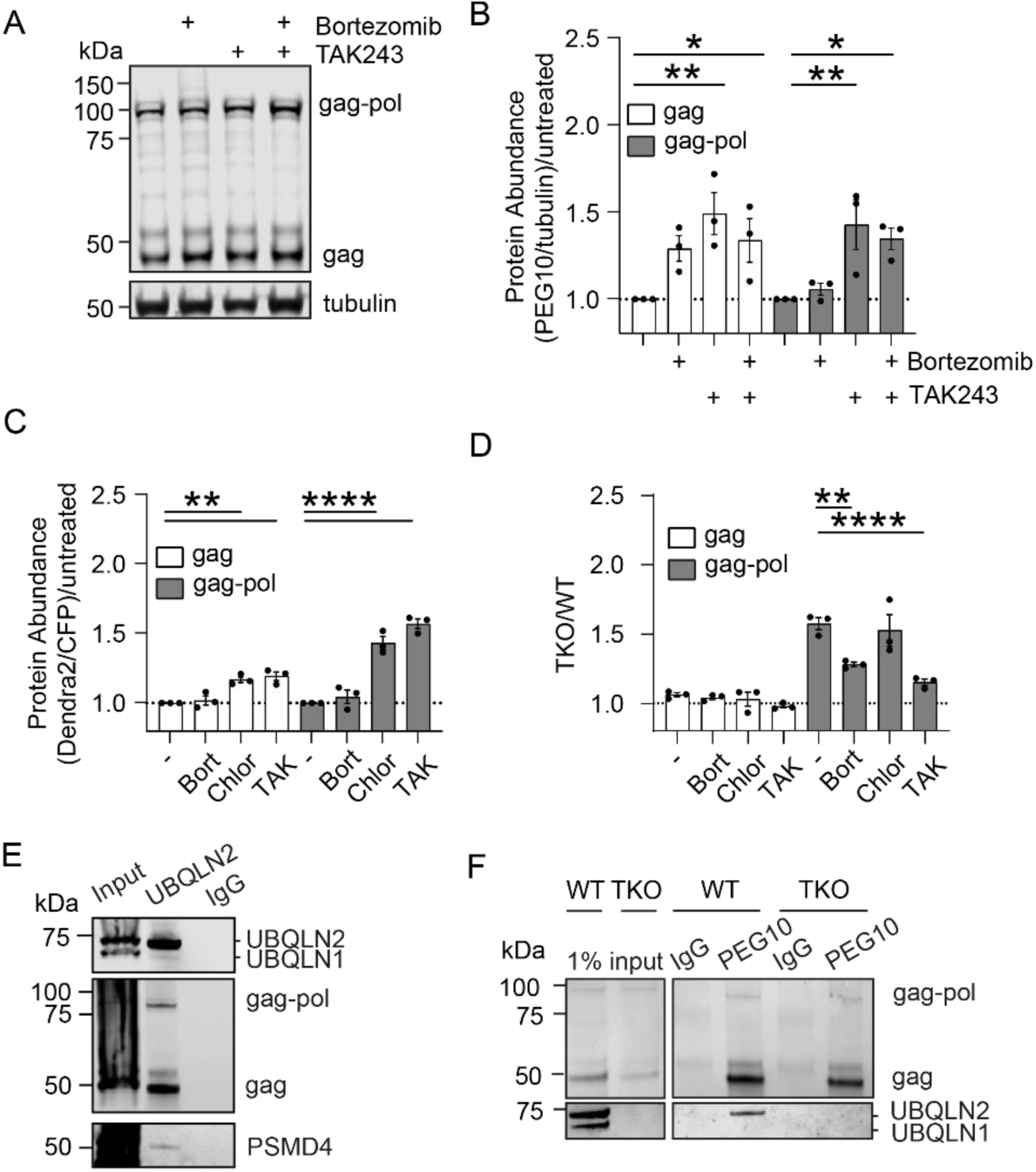
Ubiquilin-mediated degradation of PEG10 gag-pol is ubiquitin and proteasome dependent. **A)** Representative western blot of TKO HEK293 whole cell lysate. Cells were treated with drug for 16 hours prior to harvest. PEG10 was probed with polyclonal PEG10 antibody. Shown is one of three representative experiments. **B)** Quantification of data shown in (A), where samples were normalized to the abundance of each protein in untreated cells for each individual experiment. Statistics were determined by two-way ANOVA with multiple comparisons, n=3. **C)** Plotted data for client accumulation assay performed in TKO cells using gag and gag-pol with and without (-) drug treatment for 16 hours prior to flow cytometry. Like (B), samples were normalized to the abundance of each protein in untreated cells for each individual experiment. Statistics were determined by one-way ANOVA with multiple comparisons, n=3. **D)** TKO/WT values were generated from **Figures 2C and 1E** by dividing TKO values by WT values for each paired experiment. Statistics were determined by two-way ANOVA with multiple comparisons, n=3. **E)** Immunoprecipitation of endogenous UBQLN2 using UBQLN2 specific antibody or IgG control from WT cells followed by western blot analysis shows co-precipitation of PEG10 gag and gag-pol, as well as the proteasomal subunit PSMD4. The UBQLN1/2 antibody used for western blot recognizes both Ubiquilins. Shown is one of two representative experiments. **F)** WT and TKO HEK293 cells were subjected to an immunoprecipitation of endogenous PEG10 using PEG10 polyclonal antibody or IgG control followed by western blot analysis. Western blot was probed for PEG10 with polyclonal antibody and UBQLN1/2 using an antibody that recognizes both Ubiquilins. Shown is one of two representative experiments.

To summarize our findings, paired data from **Figures 1E** and **2C** of transfected WT and TKO cells were transformed to obtain TKO/WT values, where Ubiquilin clients that accumulate in TKO cells have values >1, whereas values ≤1 represent non-clients. Gag showed no regulation by Ubiquilins regardless of drug treatment (**Figure 2D**, **left**), as evidenced by the values of ∼1 in all culture conditions. Without drug treatment, gag-pol levels were higher in TKO cells, which was mitigated by inhibition of the proteasome or ubiquitination (**Figure 2D**, **right**). In contrast, treatment with chloroquine did not diminish the contribution of Ubiquilins to gag-pol abundance, indicating that autophagy and lysosomal degradation of PEG10 are not significantly impacted by the presence or absence of Ubiquilins.

The classical model of Ubiquilin-mediated protein degradation involves an interaction between the Ubiquilin and client protein to facilitate proteasomal delivery. IP of UBQLN2 in the presence of the crosslinking reagent DSP resulted in the co-precipitation of both PEG10 gag and gag-pol (**Figure 2E**), consistent with previous reports (Mohan *et al*., 2024). PSMD4 (also known as Rpn10), a subunit of the proteasome and a known UBQLN2 interactor (Ko *et al*., 2004; Hamazaki, Hirayama and Murata, 2015), was also visible (**Figure 2E**). IP of endogenous PEG10 demonstrated a reciprocal co-precipitation of UBQLN2 (**Figure 2F**). Notably, while UBQLN2 and UBQLN1 are known to interact (Ford and Monteiro, 2006; Lee and Brown, 2012), minimal UBQLN1 signal was visible in the eluate of the UBQLN2 or PEG10 IP (**Figure 2E-F**), highlighting the selectivity of the interaction in this system.

To exclude an indirect contribution of other Ubiquilins to a UBQLN2:PEG10 interaction, a TKO cell line with doxycycline inducible expression of MYC-UBQLN2 was used for IP. As before, IP of MYC-UBQLN2 co-precipitated both gag and gag-pol protein, and IP of PEG10 resulted in co-precipitation of MYC-UBQLN2 (**Supplementary Figure 2C**). ALS-associated UBQLN2 mutations of P497H, which has a minor effect on PEG10 abundance, and P506T, which has a more significant effect on PEG10 abundance (Black *et al*., 2023), did not visibly reduce binding capacity to PEG10 (**Supplementary Figure 2D**), though it remains possible there are minor changes to affinity which are not readily apparent in these tests.

### Lysine residues of the pol region of PEG10 are critical for UBQLN2-dependent proteasomal degradation

PEG10 contains fifteen lysine residues in its gag region, one lysine in the frameshift region, and four (393, 567,586, 590) in its pol region (**Figure 3A**), which could be ubiquitinated to direct protein degradation. Previous work shows many of these residues are ubiquitinated in human cell lines (Hornbeck *et al*., 2015; Abed *et al*., 2019), but it is not known how each lysine influences protein stability. By mutating individual lysine residues to arginine (K>R), the contribution of specific residues to UPS-mediated degradation can be directly interrogated. Because of the unique UBQLN-dependence of gag-pol protein, lysines in the pol region of PEG10 were subjected to mutation first. WT and TKO cells were transfected with K>R mutant reporter constructs and protein abundance was plotted for each as TKO/WT. When all four pol lysines were mutated to arginine, gag-pol lost all dependence on UBQLNs for its degradation (**Figure 3B**, **3D**). However, no single K>R mutations of pol lysines had any significant effect on UBQLN-dependent accumulation (**Figure 3B**). When two K>R mutations were combined, they had variable effects, suggesting a dominant role of K567 in regulating UBQLN-mediated degradation of PEG10 (**Figure 3C-D**). When three residues were mutated K>R, dependence on UBQLNs was also completely lost (**Figure 3C-D**), implying a negligible or secondary role of lysine 393 in UBQLN-mediated targeting. Consistent with a role for UBQLNs in facilitating degradation by the ubiquitin-proteasome pathway, PEG10 gag-pol^K>R^ did not meaningfully accumulate upon proteasome inhibition, but did accumulate upon autophagy or E1 inhibition (**Figure 3E**), although WT gag-pol protein had a much larger accumulation upon E1 inhibition in comparison (**Figure 1E**). Together, they suggest that mutation of pol region lysine residues significantly impacts Ubiquilin-dependent degradation via the ubiquitin-proteasome system.

**Figure 3:**
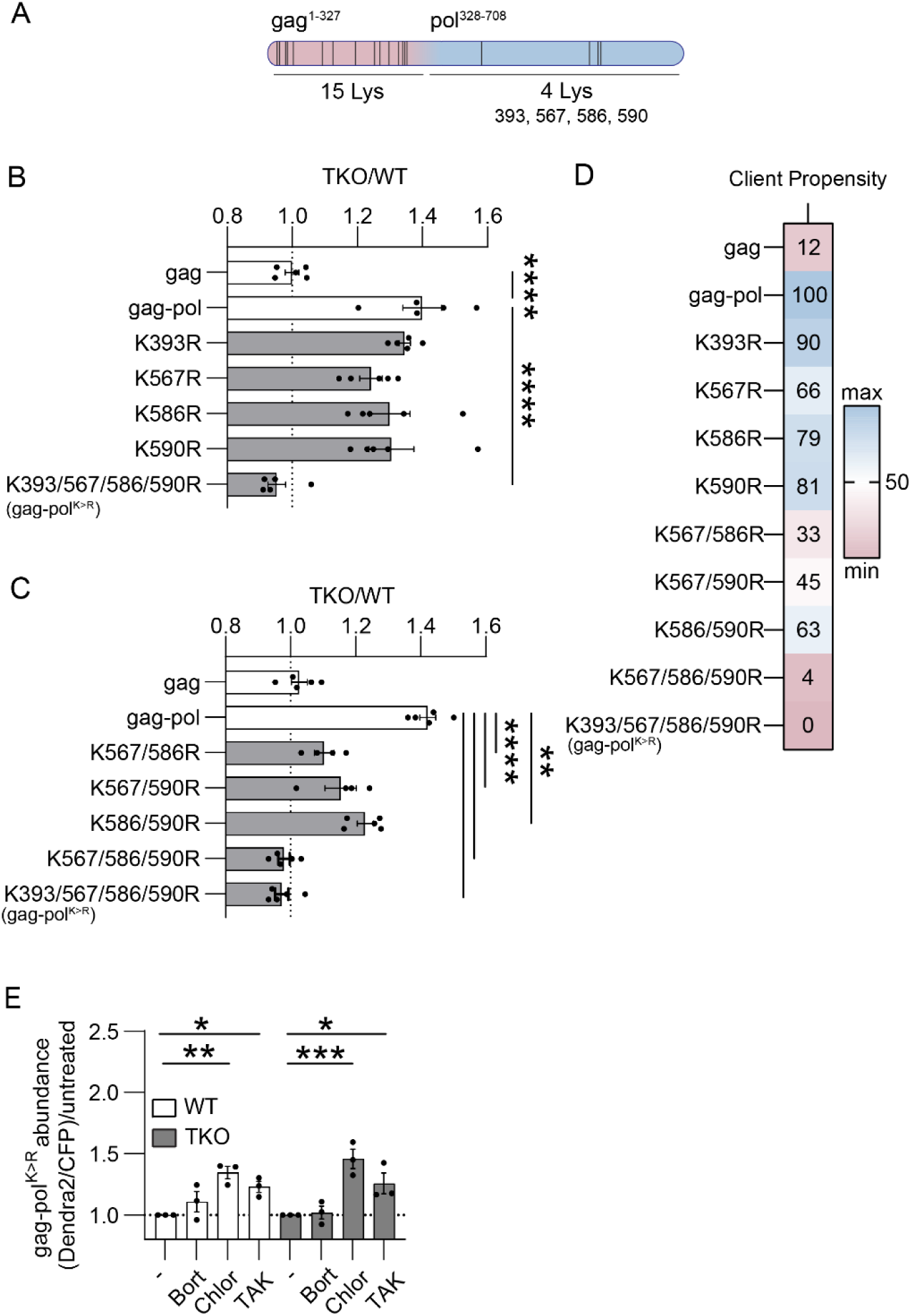
PEG10 pol lysine residues are necessary for UBQLN-mediated degradation. **A)** General schematic showing the approximate location of lysine residues along PEG10 gag-pol. **B)** Plotted cumulative data for client accumulation assay of single K>R mutations, and **C)** combinatorial K>R mutations. All experimental data is shown, but statistics were calculated by one-way ANOVA with multiple comparisons using gag-pol as a control and only paired samples within an experiment. **D)** Client propensity heatmap on the right was calculated by averaging all TKO/WT values for the experiments in B and C, normalized to the minimum and maximum of the dataset. **E)** K393/567/586/590R, or gag-pol^K>R^, was subjected to the client accumulation assay. Cells were treated with or without (-) the drugs shown for 16 hours prior to flow cytometry. Samples were normalized to the abundance of each protein in untreated cells for each individual experiment. Statistics were determined by one-way-ANOVA with multiple comparisons, n=3.

To determine the contribution of lysines in the gag region to PEG10 degradation, all lysine residues within gag were mutated to arginine. In WT cells, K>R mutations resulted in increased Dendra2 abundance, which was not observed in TKO cells (**Supplementary Figure 3A**). As expected, gag^K>R^ was not a client of Ubiquilins (**Supplementary Figure 3**); however, gag^K>R^-pol was also not a client of Ubiquilins (**Supplementary Figure 3**), indicating that at least one lysine residue in the gag region is also required for Ubiquilin-mediated degradation.

### PEG10 binding to UBQLN2 is UBA domain, lysine, and ubiquitin independent

Ubiquilin:client binding is thought to be governed by STI1:protein interactions and UBA:ubiquitin chain interactions (Lee and Brown, 2012; Zheng, Yang and Castañeda, 2020). Canonically, binding to ubiquitinated substrates is ablated by deletion of the UBA (ΔUBA), while proteasomal delivery is impaired upon deletion of the UBL (ΔUBL) (Osaka, Ito and Suzuki, 2016). To test the contribution of these domains to PEG10 binding, TKO HEK293 cells were transfected with FLAG-UBQLN2 constructs (Gerson *et al*., 2020) – WT, ΔUBL, and ΔUBA – and lysate was subjected to a FLAG-IP. Endogenous PEG10 gag and gag-pol reactive bands were visible in all conditions, regardless of UBQLN2 deletion (**Figure 4A**), indicating that the UBL and UBA domains are unnecessary for client interaction.

**Figure 4:**
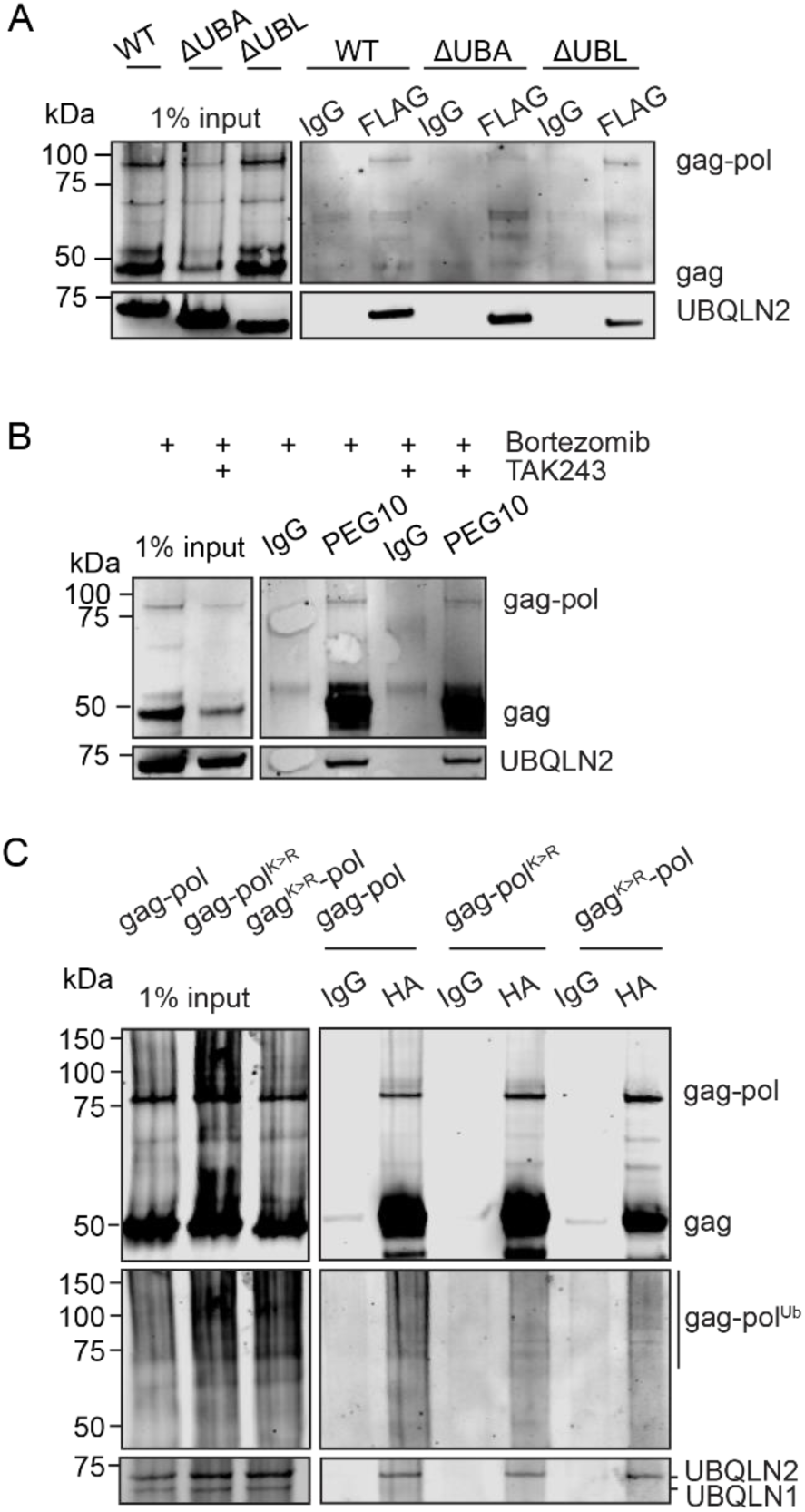
UBQLN2 binds to PEG10 in a ubiquitin-independent manner. **A)** TKO HEK293 cells were transiently transfected with N-term 3xFLAG-UBQLN2 constructs and lysate was subjected to an immunoprecipitation using an anti-FLAG antibody or IgG control antibody followed by western blot. PEG10 was detected with polyclonal antibody, and UBQLN2 was detected with FLAG antibody. Shown is one of two representative experiments. **B)** MYC-UBQLN2-expressing cells were drug treated for 16 hours, and lysate was subjected to an immunoprecipitation using a PEG10 or IgG control antibody followed by western blot analysis. PEG10 was detected with polyclonal antibody and UBQLN2 was detected by anti-MYC-tag antibody. Shown is one of two representative experiments. **C)** WT HEK293 cells were transiently transfected with HA-tagged PEG10 constructs and a plasmid expressing 3xFLAG-Ub. An immunoprecipitation was performed via either HA-tag antibody or IgG control and samples were visualized by western blot. PEG10 was probed with HA antibody, Ubiquitin was probed with FLAG antibody, and UBQLN2 was visualized with an antibody that recognizes both UBQLN1 and UBQLN2. Shown is one of two representative experiments.

Binding of PEG10 to UBQLN2^ΔUBA^ implied that ubiquitination of PEG10 was dispensable for their interaction. To more directly test the role of ubiquitination in UBQLN2-PEG10 binding, MYC-UBQLN2 expressing cells were treated with TAK243 for an endogenous PEG10 IP. A MYC-UBQLN2 band was present regardless of TAK243 treatment (**Figure 4B**), indicating that ubiquitination is not necessary for the interaction between UBQLN2 and PEG10.

Finally, cells were transfected with HA-tagged K>R mutants and 3xFLAG-Ub, followed by an IP of HA-tagged PEG10. These constructs still co-precipitated UBQLN2, indicating that lysine-mediated ubiquitination of PEG10 is not necessary for interaction with UBQLN2 (**Figure 4C**). Therefore, while ubiquitination is necessary for gag-pol degradation, it is dispensable for UBQLN2 binding.

### UBE3A, an E3 ligase and binding partner of UBQLN2, alters gag-pol ubiquitination and abundance

While ubiquitination of PEG10 is dispensable for its interaction with UBQLN2, it is still necessary for UBQLN-dependent, proteasomal degradation (**Figure 1**, **Figure 3**). Furthermore, binding of gag-pol by UBQLN2 is not sufficient for degradation to occur, as gag-pol^K>R^ interacts with UBQLN2, but is not degraded by the proteasome. One model of UBQLN1-mediated degradation of mislocalized Omp25 determined that binding of Omp25 by UBQLN1 resulted in recruitment of an unknown E3 ligase to facilitate ubiquitination and subsequent degradation of the protein (Itakura *et al*., 2016). On the other hand, recent work has also shown UBQLN2-mediated protection of ATP5G1 through facilitated interaction with the E3 ligase SCF^bxo7^ (Scheutzow *et al*., 2024). Considering these findings, our results led us to examine the E3 ligase that may work with UBQLN2 to facilitate ubiquitin-mediated degradation of PEG10.

Recent work has linked the HECT E3 ligase UBE3A (also known as E6AP) to PEG10 ubiquitination and regulation in neurons with implications for the neurodevelopmental disorder Angelman’s Syndrome (Pandya *et al*., 2021). UBE3A is well-known for its oncogenic role in the proteasomal degradation of P53 upon binding to the human papillomavirus E6 protein (Yamamoto, Huibregtse and Howley, 1997); however, mutations in the *UBE3A* gene also cause the neurodevelopmental disease Angelman’s syndrome (Kishino, Lalande and Wagstaff, 1997). In addition to P53, UBE3A has been linked to the ubiquitination and degradation of many additional host proteins (Martínez-Noël *et al*., 2018) including a retrotransposon-derived gag-like protein Arc (Greer *et al*., 2010) that resembles PEG10 and plays important and unique roles in neurological function (Chowdhury *et al*., 2006; Ashley *et al*., 2018; Pastuzyn *et al*., 2018). UBE3A has also been previously linked to the Ubiquilin family (Kleijnen *et al*., 2000; Kleijnen, Alarcon and Howley, 2003), and structural work has shown the N-terminal AZUL domain of UBE3A binds to the UBA of UBQLN1 and UBQLN2 (Buel *et al*., 2023). Therefore, we sought to test the hypothesis that UBE3A contributes to the UBQLN2-dependent proteasomal degradation of PEG10.

UBE3A depletion experiments confirmed a role for the E3 ligase in regulating gag-pol abundance and revealed a dependence on UBQLNs for this control. WT and TKO HEK293 cells were transfected with non-targeting control (NTC) or *UBE3A*-targeting siRNAs (either as a pooled mixture or a single siRNA) followed by western blot for endogenous PEG10 (**Figure 5A-D**, **Supplementary Figure 4**). In WT cells, knockdown of *UBE3A* did not influence gag protein levels (**Figure 5C**, **left**) but increased gag-pol abundance significantly (**Figure 5D**, **left**). Both the 80 kDa and higher molecular weight forms of gag-pol protein were affected (**Figure 5D-E**, **left**); this unexpected result suggests there are additional E3 ligases which ubiquitinate PEG10 gag-pol, and that UBE3A regulates the abundance of these ubiquitinated forms of gag-pol, as well. In contrast, TKO HEK293 cells showed much smaller changes to gag or gag-pol abundance upon *UBE3A* knockdown (**Figure 5A-E**, **Supplementary Figure 4**), suggesting that the absence of Ubiquilins prevents UBE3A from regulating PEG10. Examination of gag-pol molecular weight showed a very minor increase to abundance of both the 80 kDa species and higher molecular weight species (**Figure 5E**), indicating the absence of Ubiquilins influences both populations.

**Figure 5:**
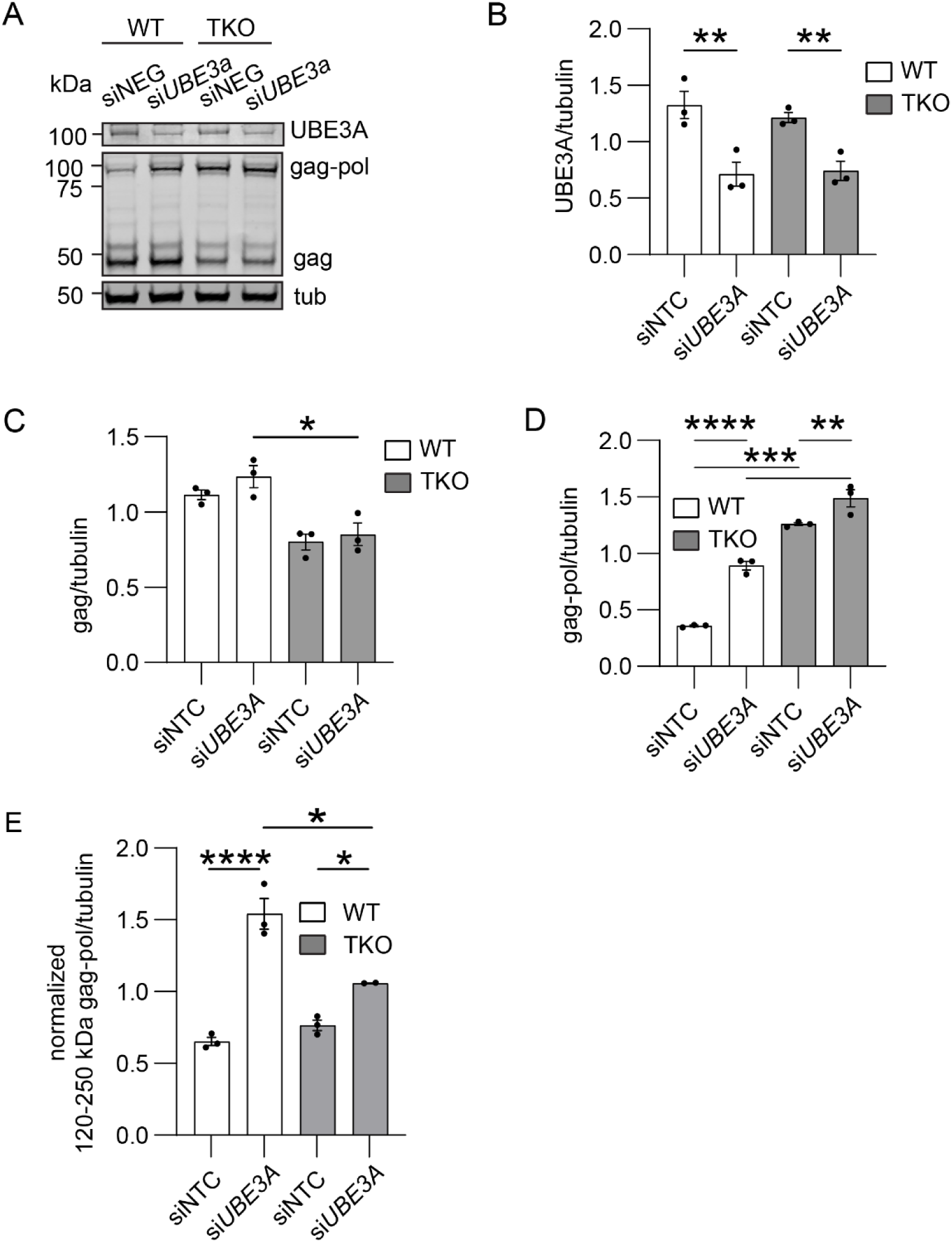
UBE3A knockdown results in accumulation of gag-pol protein. **A)** Representative western blot of *UBE3A* knockdown in WT and TKO HEK293 cells. Cells were transfected with 100 pmol siRNA against a non-targeting construct (*NTC*) or *UBE3A* and harvested for western blot 72 hours later. PEG10 and UBE3A were visualized with polyclonal antibodies. Shown is one of three representative experiments. **B)** Quantification of UBE3A protein abundance from (A). **C)** Quantification of gag protein abundance from (A). **D)** gag-pol protein abundance from (A). **E)** Quantitation of high molecular weight (120-250 kDa) gag-pol protein from (A). For (B-E), results for each band were normalized to a blot average of that band for each independent experiment, and statistics were determined by ANOVA with multiple comparisons, n=3.

IP of UBQLN2 co-precipitated UBE3A along with PEG10 and the proteasome subunit PSMD4 (**Figure 6A**). To test the contribution of the AZUL domain to this interaction, we transfected cells with FLAG-tagged UBE3A WT (UBE3A^FL^) or ΔAZUL (UBE3A^ΔAZUL^) and performed the same IP. Both constructs co-precipitated with UBQLN2 and PEG10 (**Figure 6B)**, though UBE3A^ΔAZUL^ may have a partial defect in UBQLN2 binding. Despite the slight decrease in UBE3A intensity in the ΔAZUL condition, PEG10 proteins were co-precipitated to equal amounts with UBQLN2 (**Figure 6B**), suggesting that PEG10 binding to UBQLN2 is independent of UBE3A. In conclusion, UBE3A associates with UBQLN2 and PEG10, and in this system, its interaction is not entirely dependent on the presence of the AZUL domain.

**Figure 6:**
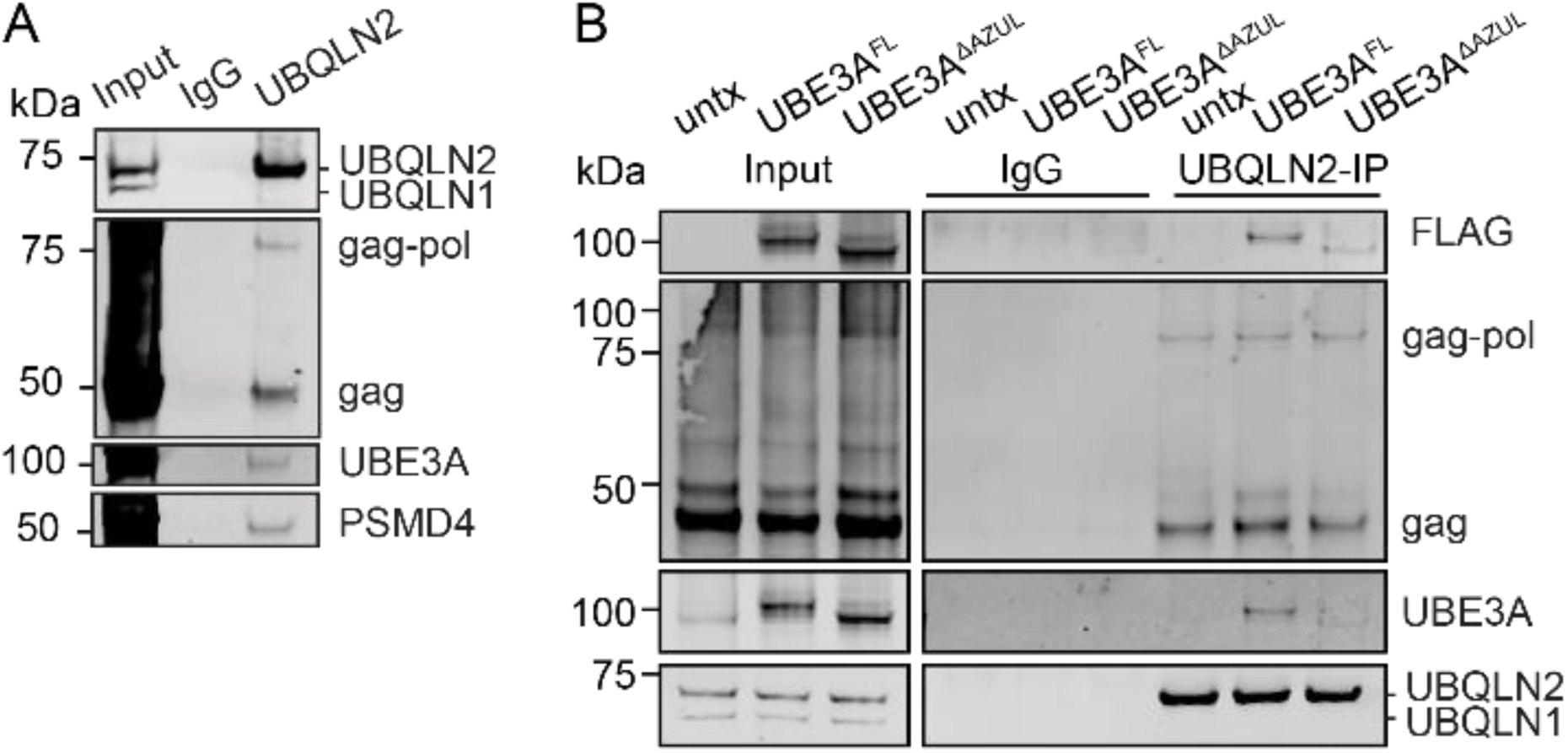
UBE3A binds to UBQLN2 and PEG10. **A)** WT HEK293 cell lysate was subjected to an immunoprecipitation with anti-UBQLN2 antibody or IgG control and detected using western blot. UBQLN1/2 were detected using an antibody that recognizes both Ubiquilins. **B)** WT HEK293 were transfected with 100 pmol siRNA against a non-targeting construct (NTC) or UBE3A and 24 hours later co-transfected with HA-gag-pol. After 48 hours, cells were lysed and PEG10 was immunoprecipitated by anti-HA antibody or IgG control, followed by western blot. UBE3A was detected with FLAG antibody and PEG10 was detected with polyclonal antibody. n=3.

## Discussion

Here, we have explored the stepwise pathway of UBQLN2-dependent proteasomal degradation of the retrovirus-like protein, PEG10. Two alternative PEG10 proteins are formed resulting from the infrequent utilization of a -1 ribosome frameshift: a short form, referred to as gag, which is not a client of UBQLN2, and a long form, gag-pol, which requires UBQLN2 for its degradation. One initial focus of our work was the determination of degradation pathway for the PEG10 proteins. We found gag-pol to be uniquely dependent on the proteasome for its degradation, whereas both gag and gag-pol accumulated upon inhibition of autophagy and ubiquitination (via E1 inhibition).

While it has been reported Ubiquilins can facilitate ubiquitin-independent proteasomal degradation (Makaros *et al*., 2023), the Ubiquilin-dependent degradation of gag-pol was dependent on functional ubiquitination. Immunoprecipitated PEG10 protein probed against ubiquitin showed a smear of ubiquitination at, and above the molecular weight of gag-pol protein, while gag protein demonstrated minimal ubiquitin staining. Consistent with these findings, three lysine residues (567,586,590) within the pol region were necessary for the degradation of the gag-pol protein, suggesting their ubiquitination contributes to proteasomal degradation of the protein. Ubiquitination of the pol region was not sufficient for proteasomal degradation, as a gag^K>R^-pol mutant was also defective in proteasomal degradation. We conclude that ubiquitination of both gag and pol regions are necessary but not sufficient for degradation by the proteasome, which could explain why gag is not a client despite binding to UBQLN2 (Mohan *et al*., 2024).

Probing of Ubiquilin-client interactions has shown a contribution of STI1 domains as well as UBA:ubiquitin interactions (Kurlawala *et al*., 2017). For PEG10, we found the UBA:ubiquitin interaction appeared dispensable for binding between UBQLN2 and PEG10, as inhibition of ubiquitination and deletion of the UBA domain did not impact binding. These data are consistent with recent work highlighting the importance of STI1 domains in binding mildly hydrophobic and alpha helical protein sequences (Fry, Saladi and Clemons Jr, 2021; Onwunma *et al*., 2024). This data is also in line with reports that UBQLN2 binding to PEG10 is facilitated by RTL8, a related retrotransposon of PEG10 within the sushi-ichi family of retrotransposons (Mohan *et al*., 2024). Future work is necessary to determine which regions of PEG10 gag contribute to this interaction, but these findings strengthen the model of Ubiquilin:client interaction where STI1 domains dominate client binding and suggests the UBA domain plays a role independent of client binding.

In light of these findings, it is notable that perturbation of UBQLN1 and UBQLN4 have no effect on PEG10 abundance (Black *et al*., 2023), suggesting UBQLN2 is unique in its ability to regulate PEG10. Further, despite published interactions between UBQLN1 and UBQLN2 (Chang and Monteiro, 2015), IP of PEG10 failed to co-precipitate detectable amounts of UBQLN1. Given these results, we conclude there is minimal interaction of UBQLN1 with UBQLN2 and PEG10 in this system, which could reflect the high PEG10 levels in these cells or a difference in IP methods.

One potential contributor to the unique relationship between UBQLN2 and PEG10 may be the proline-rich PXX domain of UBQLN2, which is not found in any of the other Ubiquilin proteins. Mutation of the PXX domain is frequently observed in UBQLN2-mediated ALS-FTD (Deng *et al*., 2011; Lin *et al*., 2022; Stansberry *et al*., 2024) and alter the ability for the protein to form liquid-like condensates (Zheng, Yang and Castañeda, 2020; Safren *et al*., 2024), bind to the proteasome (Chang and Monteiro, 2015), and facilitate autophagosomal protein degradation (Wu *et al*., 2020). Our previous work showed UBQLN2 mutants were still capable of partially restraining PEG10 gag-pol, with UBQLN2^P506T^ leading to the most significant degradation defects (Black *et al*., 2023). These mutants showed no obvious changes to PEG10 binding; however, our data indicates that binding to UBQLN2 is not the only metric by which UBQLN-dependent degradation should be measured.

Our data showed while UBQLN2 was able to bind non-ubiquitinated PEG10 protein, ubiquitination was necessary for proteasomal degradation. Together, these pieces of data suggest client ubiquitination is subsequent to Ubiquilin binding. Therefore, we undertook additional studies to explore the mechanism of PEG10 ubiquitination. Previous studies have shown PEG10 ubiquitination is regulated by the HECT E3 ligase UBE3A, and that UBE3A binds to UBQLN2 (Pandya *et al*., 2021; Buel *et al*., 2023). *UBE3A* knockdown correlated with an increase in PEG10 protein at both the 80 kDa molecular weight and at higher molecular weights, suggesting UBE3A regulates the degradation of PEG10, and that there are other E3 ligases capable of modifying PEG10 in a way that does not influence its degradation. Consistent with this model, ubiquitin blot of precipitated PEG10 protein from cells treated with *UBE3A* siRNA did not show a major change to the ubiquitin staining pattern.

Most notably, while we observed a specific accumulation of PEG10 gag-pol protein in WT cells upon *UBE3A* knockdown, there were comparatively minimal changes to TKO cells. This indicates UBQLN2 is necessary for the UBE3A-mediated proteasomal degradation of gag-pol protein. In the absence of UBQLN2, the 80 kDa molecular weight gag-pol protein accumulates dramatically, leading us to conclude that UBQLN2 is necessary for the UBE3A-mediated ubiquitination of PEG10. This could explain why previous attempts to observe PEG10 ubiquitination by UBE3A *in vitro* were unsuccessful (Pandya *et al*., 2021), although more work is necessary to confirm the presence of UBQLN2 as sufficient to promote direct ubiquitination of PEG10 by UBE3A.

These findings are particularly interesting in light of complementary work on the role of ubiquitin in modulating phase separation of UBQLN2. UBQLN2 can form liquid condensates (Gerson *et al*., 2021) and to localize to stress granules in living cells (Alexander *et al*., 2018), and the addition of ubiquitin causes UBQLN2 phase separations to disperse (Dao *et al*., 2018; Castaneda, Dao and Yang, 2022). These findings have also contributed to the theory that client ubiquitination is an important step in the pathway of Ubiquilin-mediated protein degradation (Dao and Castañeda, 2020; Valentino *et al*., 2024). In line with this, our model shows a multistep pathway of Ubiquilin-mediated proteasomal degradation. In the first step, UBQLN2 binds to a non-ubiquitinated PEG10, and UBE3A, in unknown order. UBE3A then ubiquitinates PEG10 gag-pol, which is necessary for its proteasomal degradation. This stands in contrast to previous models where UBA:ubiquitin interactions dominate Ubiquilin-client binding, and emphasizes the presence of a new cofactor: the E3 ligase. Our current understanding and model show a need for the interactions of multiple domains and unique residues of UBQLN2, PEG10, and UBE3A that are necessary for binding and subsequent protein degradation. While the field has sought to understand the role of Ubiquilins in protein degradation, we have shown a novel set of UBQLN2 client characteristics which may expand the search for other Ubiquilin clients. Additionally, this body of work shows the client specificity of UBQLN2, and highlights the possibility of other specific Ubiquilin, client, and E3 ligase machinery.

## Materials and Methods

### Cloning

All *Homo sapiens* PEG10 [gag (AA 1-325), gag-pol (AA 1-708)] and UBE3A isoform 2 [FL (AA 1-853) ΔAZUL (AA 60-853) ((Kleijnen *et al*., 2000)] plasmids were cloned into pCDNA3.1 or pDendra2 backbone designed with a CMV promoter, using GeneArt Gibson Assembly (Invitrogen). For PEG10 lysine to arginine mutants of the pol region, Site Directed Mutagenesis was used. All HA-tagged constructs had an N-terminal 2xHA-tag. All FLAG-tag constructs made in this study had an N-terminal 3xFLAG-tag. All Dendra2 tagged constructs had a C-terminal Dendra2 tag.

pCMV4-FLAG-UBQLN2 [WT (AA 1-624), ΔUBL (AA 1-624 missing AA 33-107), and ΔUBA (AA 1-580)] were a gift from Dr. Lisa Sharkey (Kleijnen *et al*., 2000; Mohan *et al*., 2022). PEG10 gag^K>R^ was purchased from Twist Biosciences before PCR and Gibson assembly to generate gag^K>R^-pol constructs. The 3xFLAG-Ubiquitin plasmid generated in this paper was subcloned from an HA-Ubiquitin plasmid (Addgene: 18712). WT UBE3A was amplified directly from HEK293 cDNA, then Gibson primers were designed to ensure inclusion or deletion of the AZUL domain. Chemically competent DH5α E. coli (Invitrogen) were transformed with plasmid and plated on either 50 µg/mL kanamycin (Teknova, cat#K211) or 100 µg/mL carbenicillin (Gold Biotechnology, cat #C-103-5) in LB agar (Teknova, cat #L9115) for 16-18 hours. Singles colonies were selected and sequenced with either Sanger sequencing (Azenta) or whole-plasmid sequencing (Plasmidsaurus). Sequence confirmed plasmids were midiprepped (Zymo, cat #D4201) and stored at approximately 1 µg/µL of purified DNA in DEPC-treated water in preparation for mammalian cell transfection.

### Cell culture and cell lines

Wildtype (WT) and *UBQLN 1, 2, and* 4 triple knockout (TKO) HEK293 cells, and Flp-In T-REx 293 TKO cells expressing a doxycycline (dox) inducible MYC-UBQLN2 (WT, P497H, P506T) were a gift from Dr. Ramanujan Hegde (Medical Research Council Laboratory of Molecular Biology). Flp-In HEK293 cells (Invitrogen) expressing PEG10 HA-gag-pol were developed in-house using manufacturer’s recommendations. Cells are maintained following standard cell culture protocols, at 37°C with 5% CO2, using Dulbecco’s modified Eagle’s medium (Gibco, cat #1514016) supplemented with 1% penicillin/streptomycin (Invitrogen), 1% L-glutamine (R&D Systems, Inc.), and 10% FBS (Millipore Sigma). The concentration of contaminating doxycycline in standard cell culture FBS was sufficient to induce approximately endogenous levels of expression of MYC-UBQLN2 in Flp-In cells (Black *et al*., 2023).

For drug treatments, cells were treated for 16 hours using final concentrations of 50 nM Bortezomib (EMD Millipore, 5.04314.0001), 100 µM Chloroquine (diphosphate, VWR # 22113-5G), or 1 µM TAK243 (Selleckchem, #S8341).

### Transfection

HEK293 cells were grown to 80% confluency in cell culture dishes with a variety of well sizes depending on the end-use of the cells. Using manufacturer recommendations, lipofectamine 2000 (Invitrogen, 11668027) was used at a DNA (µg) to Lipofectamine 2000 (µL) ratio of 1:2.5, diluted in 1X Opti-MEM (Gibco, 31985062).

### Western blot

Cells were plated in 6-well plates for all western blot analyses. Cells were harvested 48 hours after transfection by gentle pipetting and spun once in PBS to wash pellets. Cell pellets were lysed in 8M urea lysis buffer (8M urea, 75 mM NaCl, 50 mM Tris pH 8.5, 1 x cOmplete Mini EDTA-free protease inhibitor cocktail tablet). Lysate was centrifuged for 10 min at 21,300 x g and supernatant was collected as whole cell lysate.

Total protein in lysate was quantified by BCA (Pierce BCA Protein Assay kit, 23227) and then samples were taken to supplement with Laemmli sample buffer with βME (Sigma Aldrich) for SDS-PAGE gel electrophoresis. Proteins were separated on 4%-12% NuPage Bis-Tris pre-cast gels (Invitrogen), and transferred to either nitrocellulose (Cytiva, 10600009) or Immobilon PVDF membranes (specifically for detection of Ubiquitin) at 20V for 60 min (Invitrogen). For detecting endogenous proteins, 9 µg of total protein was used per biological sample. For detecting transfected proteins, 5 µg of total protein was used.

Membranes were blocked using LICOR Intercept TBS or PBS blocking buffer for 30 minutes, washed three times for five minutes each using 1x TBS-T, and detected using the LICOR IR secondary antibody system. Primary antibodies for PEG10 (Proteintech, 1:1000), HA-tag (either Proteintech or Sigma, 1:5000), UBQLN2/1 (Abnova, 1:1000), M2 FLAG-tag (Sigma, 1:5000), UBE3A (Proteintech, 1:1000), PSMD4 (Cell Signaling Technology, 1:1000), MYC tag (Sigma, 1:2000), and tubulin (Novus Biologics, 1:10,000) were left on overnight at 4°C, while secondary antibodies (IR680 and IR800 conjugates, LICOR) were incubated for 1 hour at room temperature at 1:10,000 concentration. Protein bands were visualized using the LICOR Odyssey CLx and image analysis performed using LICOR ImageStudio Software.

### Flow cytometry – client accumulation assay

Cells were transfected in 48-well plates and harvested 48 hours after transfection by pipette mixing using 200 µL Flow Buffer (D-PBS, 2% FBS, 0.1% Sodium Azide). Samples were transferred into U-bottom 96-well plates and analyzed using a BD FACSCelesta fitted with a high throughput sampler with the following settings: Sample Flow Rate (1.0 µL/sec), Sample Volume (100 µL), Mixing Volume (100 µL), Mixing Speed (200 µL/sec), Number of Mixes (2), Wash Volume (200 µL). All experimental data was analyzed using FlowJo.

#### Gating strategy

Transfected cells were initially gated in FSC-A vs SSC-A with the polygon gating tool to identify ‘cells’. Within this ‘cells’ population, CFP-positive cells were gated in 405 nm vs SSC-A. A novel parameter to derive Dendra2 Green/CFP was created by dividing 488 references by 405 references, with a logarithmic scale, min= 0.0001 and max = 10. This custom parameter was utilized only in CFP positive cells, and the geometric mean was exported and used to generate all graphs.

### Immunoprecipitation (IP) and Co-IP

Antibodies (µg) against PEG10 (Proteintech), HA-tag (Proteintech), UBQLN2/1 (Abnova), or M2 FLAG-tag (Sigma) were conjugated to protein A or G resin beads (50% slurry, µL) at a ratio of 1:8, using 40 µL of resin and 5 µg of antibody per IP condition. Antibodies were incubated with the beads for 30 – 60 min at room temperature, before buffer exchange into 0.2 M Sodium Borate (pH 9.0). Antibodies were crosslinked to the beads using 20 mM DMP diluted in 0.2 M Sodium Borate (pH 9.0) for 30 minutes at room temperature with rocking. The volume of DMP solution used was 10X the resin bed volume (40 µL 50% slurry = 400 µL DMP solution). The crosslinking reaction was terminated using 0.2 M ethanolamine (pH 8.0, which was left overnight at 4°C. The next day, the resin was washed three times in 1x D-PBS before adding lysate. Resin beads were spun down at 3000 x g, for 3 minutes.

For most Co-IPs, cells were plated in 10-cm dishes and treated with bortezomib for 16 hours to induce accumulation of protein products. An exception is that endogenous UBQLN2 IPs in **Figures 2E** and **5A** were not subjected to drug treatment prior to lysis. Cells were lysed in 500 µL triton lysis buffer (200 mM NaCl, 10 mM HEPES, 10 mM EGTA, 10 mM EDTA, 1% Triton X-100, 1x cOmplete EDTA-free protease inhibitor), and centrifuged at 16,000 x g for 5 min at 4°C. Supernatant was collected and equally split between IgG and antibody-conjugated resin and incubated at 4°C overnight with an end-over-end rotator. The next day, the resin was washed three times for two minutes each, using IP Wash Buffer (1x D-PBS, % Triton X-100, 1x cOmplete EDTA-free protease inhibitor). Protein elution was achieved by the addition of IP Elution Buffer (2x Laemmli in 1x PBS), and boiling the resin for 15 minutes at 95°C. Centrifugation was used to pellet the resin, and eluate was collected before being visualized using western blot.

#### DSP Crosslinking for Co-IPs

For Co-IP experiments, cells were treated with DSP (Lomant’s Reagent, Thermo Fisher) in one of two manners (Akaki, Mino and Takeuchi, 2022). Cells were collected in tubes at 300 x g for 3 minutes and washed in 1x D-PBS. Right before use, 50 mM DSP was dissolved in DMSO, and then further diluted in 1x D-PBS for a final concentration of 1 mM. Crosslinking was performed for 30 minutes at room temperature with rocking. DSP-mediated crosslinking was stopped using DSP stop buffer (20 mM Tris pH 7.5 in PBS) for 15 minutes at room temperature with rocking. Cells were washed three times with 1x D-PBS, before lysis as previously described for IP. Alternatively, cells were washed and treated with DSP directly in 10-cm dishes before harvest and lysis prior to Co-IP.

## Statistical analysis

Statistical analyses were performed utilizing the integrated analysis suite in GraphPad Prism version 10.4.1 for Windows, (GraphPad Software). Analyses used include standard one-way ANOVA followed by Dunnett’s multiple comparisons test, and two-way ANOVA followed by Šídák’s multiple comparisons test. Statistical tests used are listed in all figure legends, error bars represent mean ± SEM, and *p<0.05, **p<0.01, ***p<0.001, and ****p<0.0001.

## Acknowledgements

We would like to thank Autumn M. Matthews for her excellent work in making and validating the stably expressed PEG10 HA-gag-pol Flp-In HEK293 cells. The authors would also like to acknowledge the Department of Biochemistry at CU Boulder: Cell Culture Facility (RRID: SCR_018988 (CCF)), the Flow Cytometry Shared Facility (RRID: SCR_019309, supported by NIH grant S10OD021601 (FLOW)), and the Share Instruments Pool, (RRID: SCR_018986).

---

**Supplemental Figure 1:**
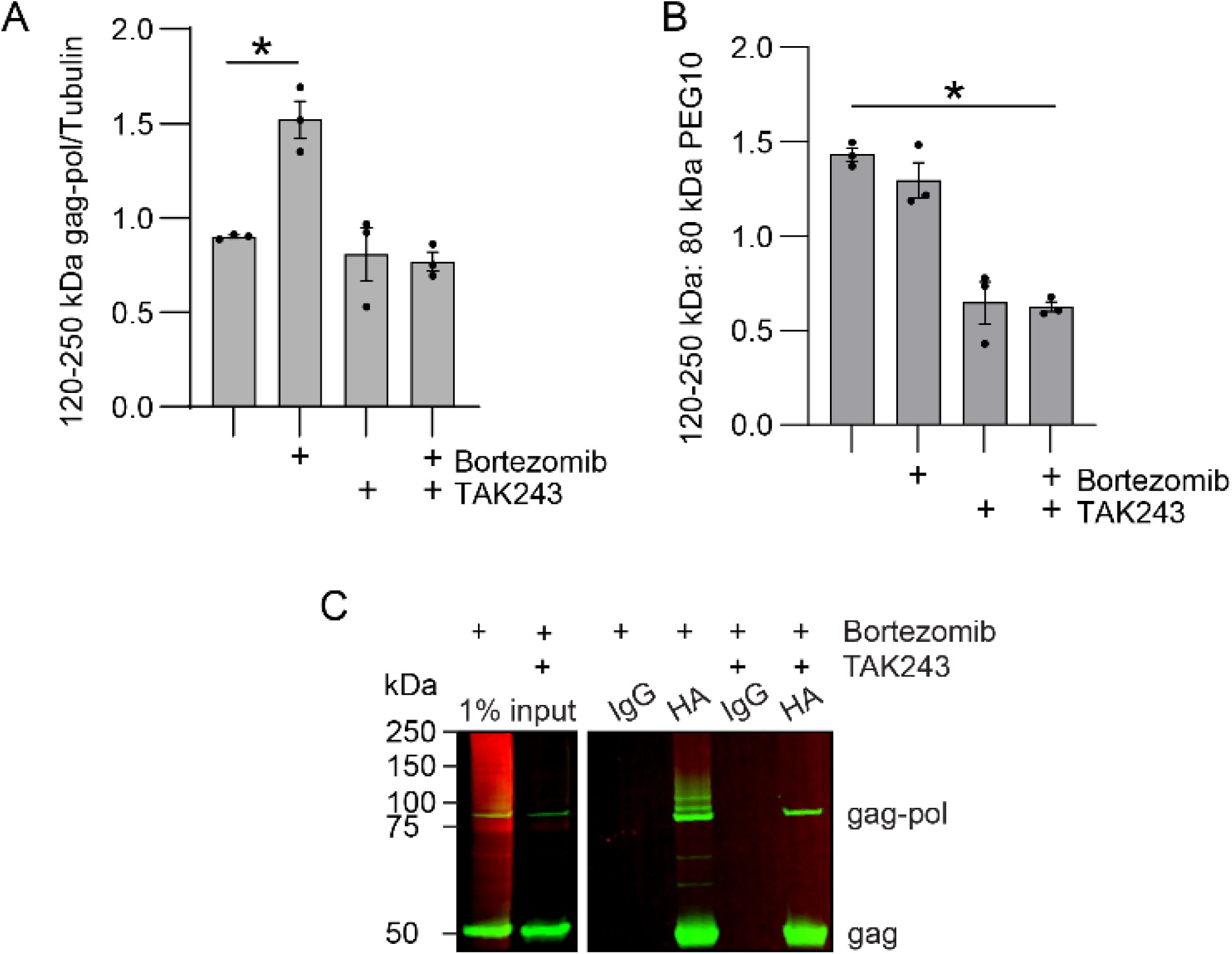
High molecular weight gag-pol is ubiquitinated protein. **A)** Quantitation of high molecular weight (120-250 kDa) gag-pol protein from Figure 1A. **B)** Ratio of high molecular weight: 80 kDa gag-pol protein for each condition tested. For (A-B), samples are normalized to a blot average to account for variance in western blot detection. Shown is mean ± SEM for three independent experiments. Statistics were determined using one-way ANOVA and multiple comparisons test. **C)** Unseparated imaging channels for Figure 1C, HA-PEG10 is in green while FLAG-Ub is in red.

**Supplemental Figure 2:**
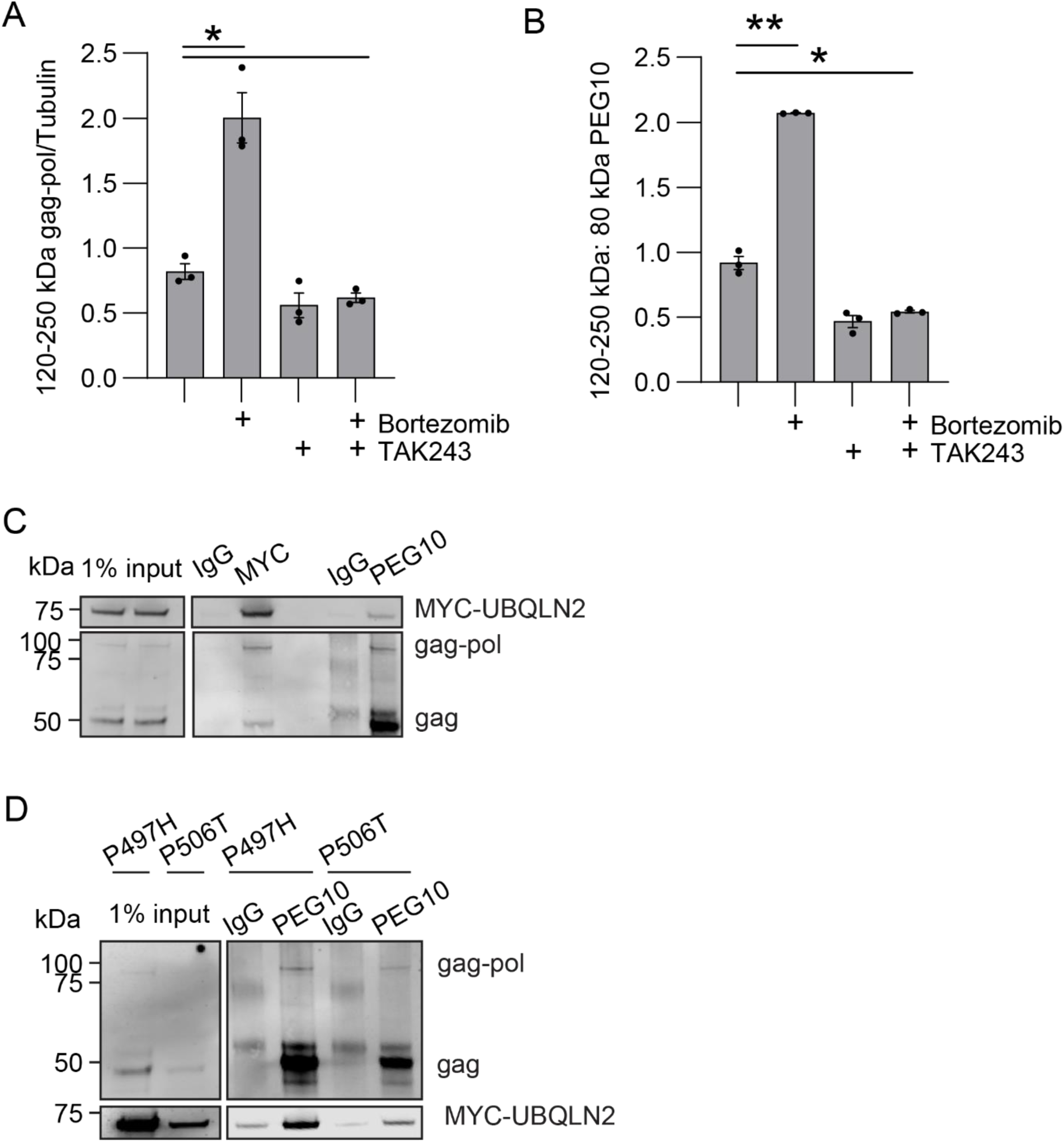
Ubiquilin deficient cells specifically accumulate 80 kDa gag-pol protein. **A)** Quantitation of high molecular weight (120-250 kDa) gag-pol protein from Figure 2A. **B)** Ratio of high molecular weight: 80 kDa gag-pol protein for each condition tested. For (A-B), samples are normalized to a blot average to account for variance in western blot detection. Shown is mean ± SEM for three independent experiments. Statistics were determined using one-way ANOVA and multiple comparisons test. **C)** Reciprocal immunoprecipitation of MYC-UBQLN2 or endogenous PEG10 shown on the same blot. Shown is one of two representative experiments. **D)** Mutant MYC-UBQLN2-expressing HEK293 cell lysate was subjected to an immunoprecipitation of endogenous PEG10, followed by western blot analysis. UBQLN2 mutants were detected by an N-term MYC-tag. Shown is one of two representative experiments.

**Supplemental Figure 3:**
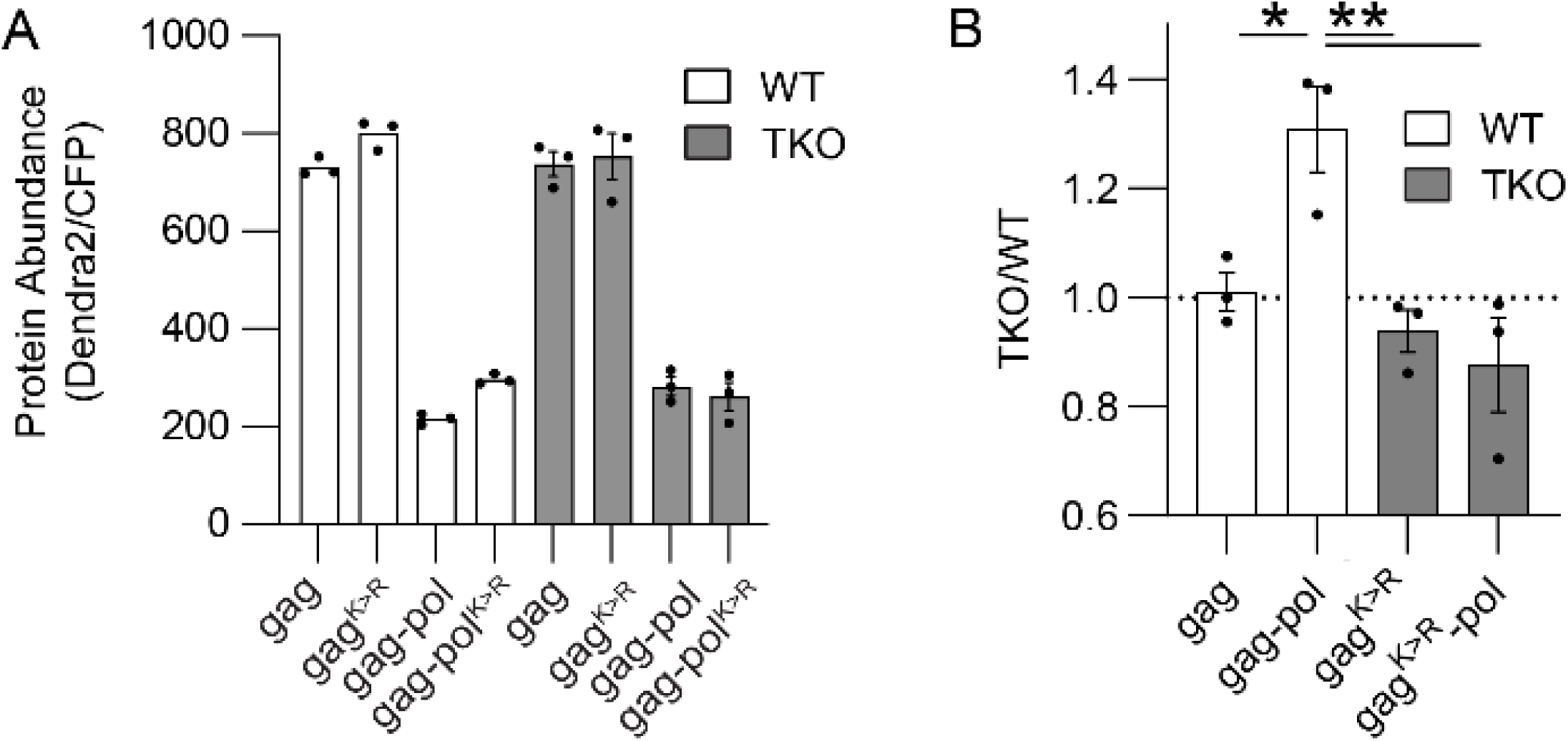
Mutation of gag lysine residues impacts protein abundance and dependence on Ubiquilins. **A)** A gag construct with all lysines mutated to arginine was cloned and used to test the importance of gag lysine residues to protein degradation. The client accumulation assay was performed in WT (left), and TKO (right) HEK293 cells. **B)** TKO /WT values are shown. For (A-B), n=3 and statistics were determined via one-way ANOVA and multiple comparisons test.

**Supplemental Figure 4:**
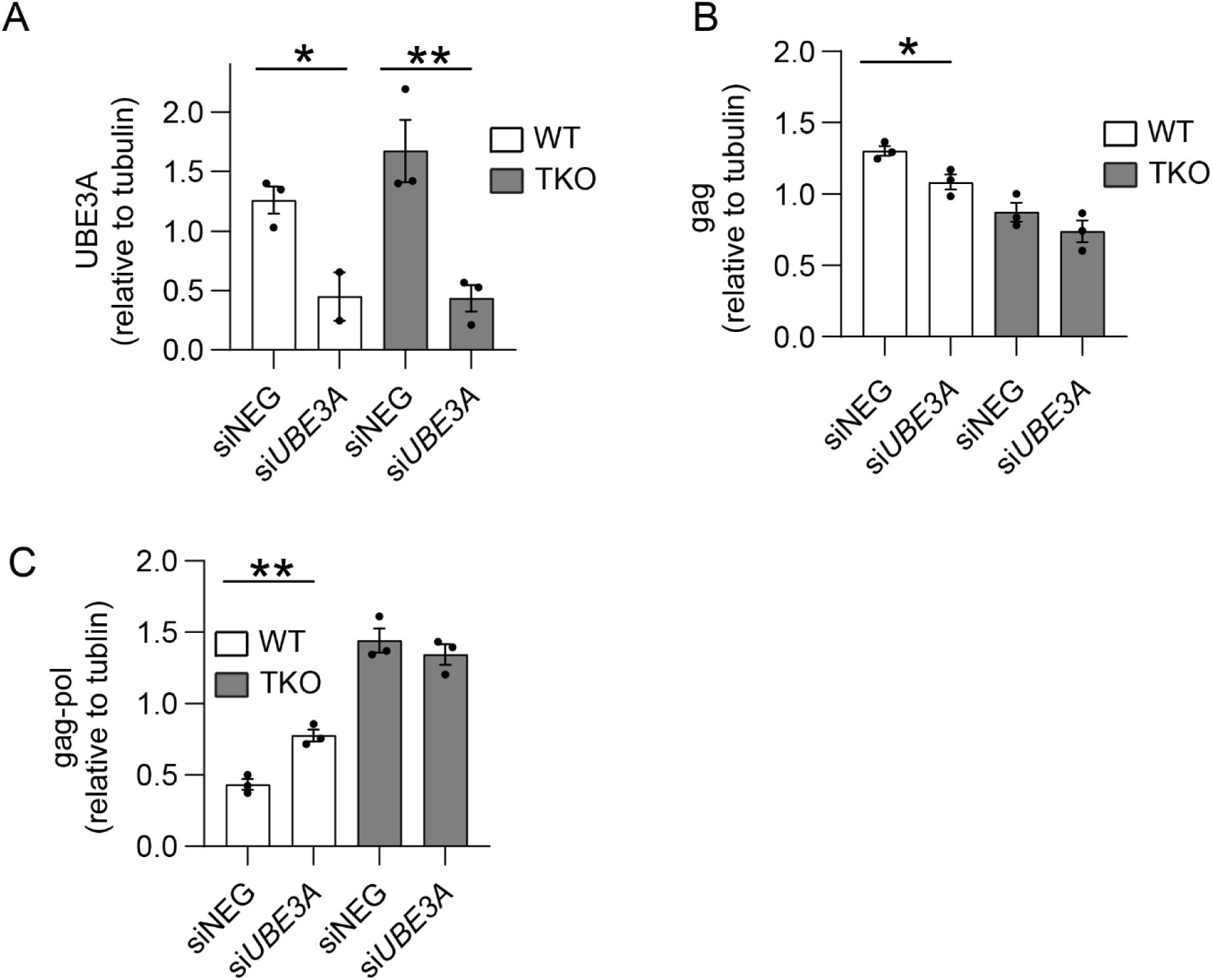
Single targeting siRNA against *UBE3A* also leads to PEG10 gag-pol accumulation only when UBQLN2 is present. **(A-C)** Quantification of western blot analysis from single si*UBE3A* siRNA in WT and TKO HEK293 cells. UBE3A (A), gag (B), and gag-pol (C) were quantitated. For (A-C), samples are normalized to a blot average to account for variance in western blot detection. Shown is mean ± SEM for three independent experiments. Statistics were determined using one-way ANOVA and multiple comparisons test.

